# Preoperative Brain Mapping Predicts Language Outcomes after Eloquent Tumor Resection

**DOI:** 10.1101/2024.05.06.592752

**Authors:** Matthew Muir, Kyle Noll, Sarah Prinsloo, Hayley Michener, Jeffrey I. Traylor, Vinodh A. Kumar, Chibawanye I. Ene, Sherise Ferguson, Jeffrey S. Weinberg, Frederick Lang, Brian A. Taylor, Stephanie J. Forkel, Sujit S. Prabhu

## Abstract

**Introduction:** Glioma patients with tumors near critical language regions present significant clinical challenges. Surgeons often lack the tools to understand how each unique surgical approach may impact linguistic ability, leading to subjective decisions and unpredictable outcomes.

**Objective:** We aim to develop an approach that uses data-driven preoperative brain mapping to quantitatively predict the impact of planned resections on long-term language function.

**Methods:** This study included 79 consecutive patients undergoing resection of language-eloquent gliomas. Patients underwent preoperative navigated transcranial magnetic stimulation (nTMS) language mapping to identify language-positive sites (“TMS points”) and their associated white matter tracts (“TMS tracts”) as well as formal language evaluations pre and postoperatively. The resection of regions identified by preoperative mapping was correlated with persistent postoperative language deficits (PLDs).

**Results:** The resection of TMS points did not predict PLDs. However, a TMS point subgroup defined by white matter connectivity significantly predicted PLDs (OR=8.74, p<.01) and exhibited a canonical group-level anatomical distribution of cortical language sites. TMS-derived tracts recapitulated normative group-level patterns of white matter connectivity defined by the Human Connectome Project (HCP). Subcortical resection of TMS tracts predicted PLDs independently of cortical resection (OR=60, p<.001). The resected TMS tract segments in patients with PLDs co-localized with normative, language-associated subcortical pathways, in contrast to the resected TMS tract segments in non-aphasic patients (p<.05). Accordingly, resecting patient-specific co-localizations between TMS tracts and normative tracts in native space predicted PLDs with an accuracy of 94% (OR=134, p<.001). Co-localization between individualized and normative tracts precisely predicted the linguistic performance of a patient intraoperatively in response to direct electrophysiological stimulation of subcortical brain.

**Conclusion:** This study outlines a data-driven brain mapping approach that provides surgical insight by preoperatively predicting the impact of individual glioma resection on long-term language function.

**Key Points:**

1. White matter connectivity determines the long-term functionality of cortical language sites mapped by TMS.
2. Long-term deficits in language processing result from resecting individualized subcortical regions within language-associated white matter tracts.
3. Non-invasive TMS language mapping combined with routine preoperative imaging can predict language outcomes of individual surgical approaches with an accuracy of 94%.

## Introduction

Gliomas account for approximately 80% of primary malignant brain tumors.^1^ Given the strong relationship between the extent of resection and survival, surgical resection remains the primary first-line treatment for glioma patients.^2,3^ These infiltrative lesions present significant challenges in surgical management due to their aggressive nature and propensity to invade critical brain regions.^1^ Operating on tumors located in the perisylvian regions of the dominant hemisphere can cause persistent postoperative language deficits (PLDs) that significantly decrease both quality-of-life and survival.^4^ Accordingly, surgeons must aim for maximal safe resection: removing as much of the tumor as possible without compromising language function.

In these patients, cortical morphometry is insufficient to guide surgical decisions due to the significant inter-individual variability of linguistic functional anatomy, both pre-existing and tumor-induced.^5–8^ Functional magnetic resonance imaging (fMRI) has been used to preoperatively localize patient-specific language regions to inform surgical decisions.^9^ However, fMRI remains limited in predicting the impact of surgical interventions on language function given its intrinsic inability to distinguish the critical versus participatory elements of the language network.^10,11^ Given the current limitations of non-invasive imaging, awake surgery with language monitoring is often chosen for patients with tumors that threaten language function.

In these settings, direct electrical stimulation (DES) has emerged as a robust technique to map patient-specific functional anatomy by correlating focal inhibitory stimulation with language task performance in real-time.^12^ While the invasive nature of the method precludes its use for surgical planning, DES can inform intraoperative decisions. However, confounders (e.g., inconsistent speech arrest interpretations, variation in stimulation intensity, and probe angle) may impact the consistency and reliability of the technique.^13^ Factors such as perioperative morbidity, intraoperative pain, seizure risk, and temporal constraints prohibit comprehensive mapping on every patient. Finally, it remains unclear whether intraoperatively identified sites of transient language disruption would lead to permanent impairment if resected.^13^ These limitations illuminate the need for data-driven surgical strategies in patients with eloquent gliomas.

Navigated transcranial magnetic stimulation (TMS) has recently emerged as a promising preoperative language mapping modality.^14,15^ It relies on a similar mechanism to DES but can apply stimulation non-invasively in an outpatient setting. The advent of navigated TMS enabled the stimulation of precise anatomical regions, allowing for personalized cortical language mapping.^16^ This study aims to combine preoperative TMS language mapping with routinely available neuroimaging to predict language outcomes after surgical resections. The results shed light on the anatomical origins of aphasic surgical deficits and inform a data-driven basis for surgical decisions.

## Methods

### Patients

This retrospective study included 79 consecutive adult presurgical glioma patients admitted between July 1st 2017 and August 1st 2023. The study included all patients diagnosed with glioma in the dominant perisylvian cortex who underwent nTMS language mapping as part of our institution’s standard preoperative protocol for eloquent tumors. Table 1 shows the demographics, baseline clinical characteristics, and surgical outcomes for this cohort. Exclusion criteria were limited to contraindications to TMS mapping, such as the presence of a pacemaker or cochlear implants. As part of the standard-of-care for eloquent tumors, patients underwent an awake craniotomy with intraoperative language mapping using DES as well as formal pre- and postoperative language evaluations by a trained neuropsychologist or speech pathologist.

**Table 1:**
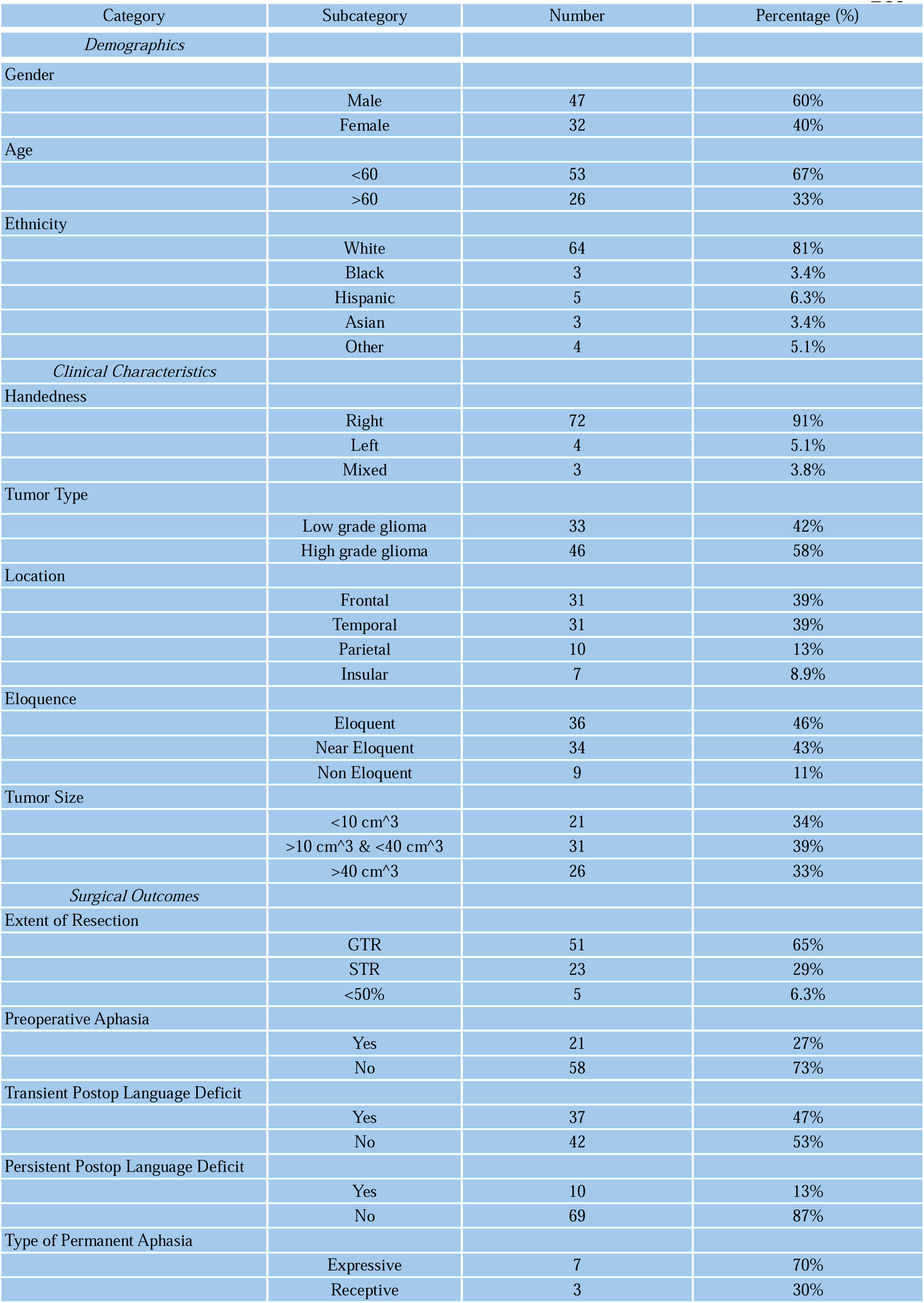
Patient demographics, clinical characteristics, and surgical outcomes.

Surgical decisions were made independently of the presurgical nTMS mapping according to the traditional standard-of-care at our institution. This study was approved by the Institutional Review Board at our institution.

### Aphasia Assessment

Patients were administered formal language evaluations by a trained neuropsychologist or speech pathologist preoperatively, immediately postoperatively, and 1-3 months postoperatively.

The formal language evaluations consisted of either the Western Aphasia Battery-Revised (WAB-R)^17^ or the following battery of neuropsychological testing: animal fluency (semantic fluency), letter fluency (phonemic fluency), and the Boston Naming Test.

A retrospective binary classification approach (new or worsened aphasia vs preserved language function) was designed to create an outcome measure that captures gross language deficits induced by the surgery. Upon immediate postoperative testing, patients with preserved language function (determined by a speech and language expert) were classified as “preserved language function” (53%) without further review of records. Patients with immediate postoperative decline (47%) were followed to the 1-3 month formal evaluation. Patients with postoperative decline at the 1-3 month evaluation (13%) were followed clinically through the 6 month mark to qualitatively verify the persistence of the deficit compared to the 1-3 month evaluation. The specific domain of impairment was derived from the 1-3 month formal evaluation compared to preoperative baseline.

For patients with an immediate postoperative deficit who also had a 1- to 3-month WAB-R assessment, the preoperative aphasia quotient (AQ) was compared to the postoperative AQ at 1 to 3 months to define the primary outcome. For those with an immediate postoperative deficit but no 1- to 3-month WAB-R assessment, the neuropsychological battery was used to classify the outcome. Given the greater sensitivity of these specific neuropsychological batteries compared to the WAB-R, an aphasic deficit was defined as a decline of at least 1.5 standard deviations from the preoperative baseline on the 1- to 3-month evaluation in at least 2 out of 3 tests.

### Presurgical Language Mapping

All patients underwent preoperative language mapping with navigated transcranial magnetic stimulation using an object naming paradigm. To define the individualized stimulation targets, an evenly spaced grid composed of stimulation points spaced 1 cm apart was overlayed onto perilesional regions with the tumour as the center point (Fig. 1B). A subset of these targets were labeled language positive (i.e., causing speech disruption) (Fig. 1C) The protocol specifics are detailed below. Correlations with intraoperative direct cortical stimulation are detailed in previous work.^14^

**Figure 1:**
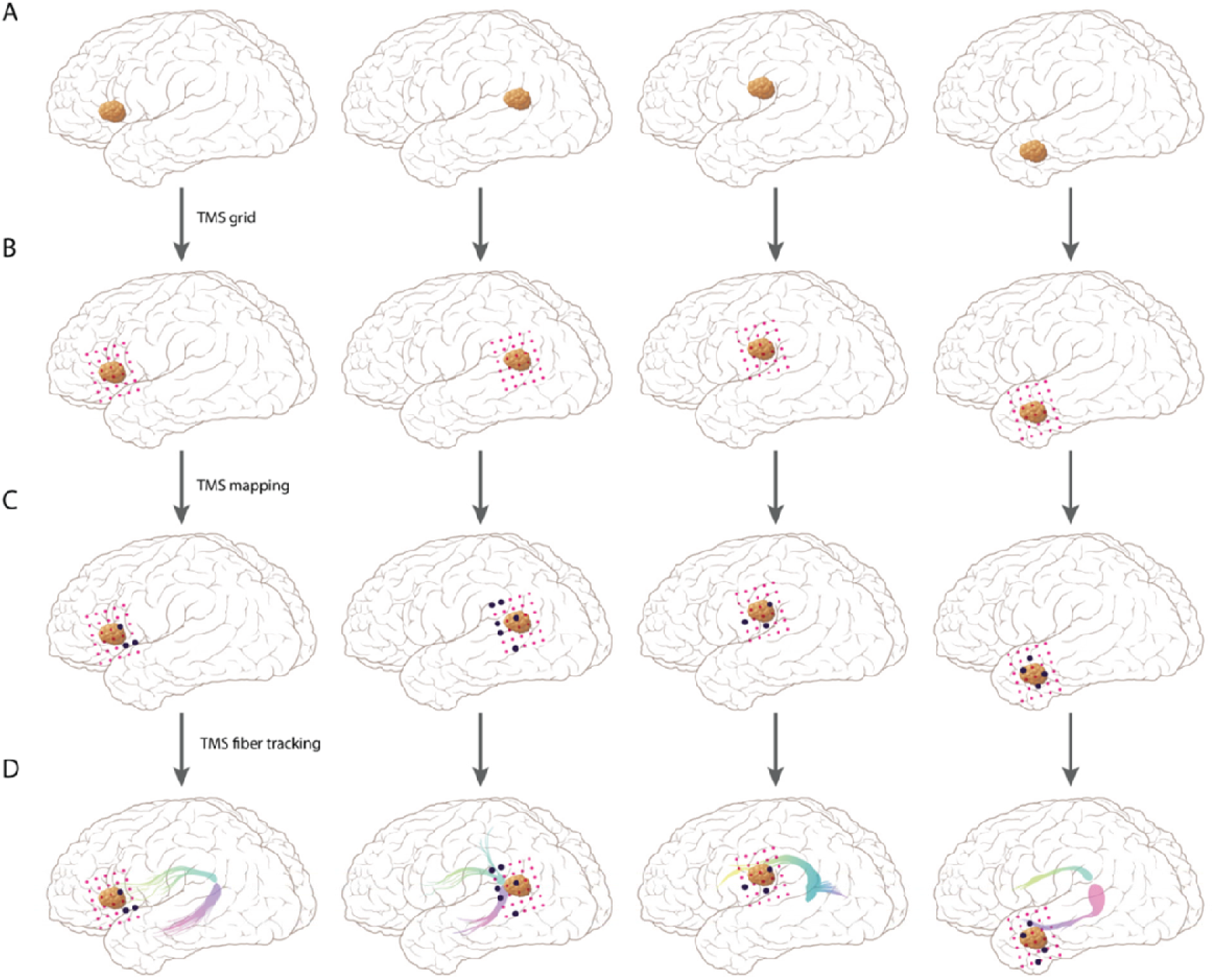
TMS fiber tracking in glioma patients. The four columns represent tumors in different locations. (B) shows placement of the grid used to guide TMS stimulation. (C) shows examples of TMS mapping results. (D) shows examples of fiber tracking based on TMS points.

Navigated TMS language mapping was performed (NBS System 3.2; Nexstim, Helsinki, Finland). TMS stimulation onset was simultaneous with presentation of line drawing objects (picture to trigger interval was 0.0 s). Time-locked rapid rate TMS trains were given for 5 Hz/5 pulses for 140 stimulations. The display time was 1,000 ms, and interpicture interval was 3,000 ms. TMS language mappings were performed according to a conventionally accepted protocol for TMS language mapping.^18,19^ The most likely location of the hand knob was identified anatomically. This area was then stimulated in a random pattern while systematically varying the rotation, tilt, and yaw of the magnetic field. The location of maximal motor evoked potential was identified. Resting motor threshold (RMT) was identified using this position.^20^ Once determined, stimulator output typically began at 110% RMT for the repetitive TMS language mapping procedure. We used a common anterior/posterior coil orientation where the coil is oriented horizontally between the nasion and external auditory meatus. Baseline object naming without stimulation was performed, which consisted of presentation of line drawings of common objects and required the patient to verbally respond with the name of the pictured object. Pictures that elicited misnaming or hesitation were discarded. Baselining was repeated until a reliable stimulus set was obtained. Target dots 1 cm apart from each other were created on the surface of the three-dimensional (3D) brain reconstruction, based on the preoperative MRI, in a grid-like pattern. One centimeter was selected because a 1 cm radius of tissue is typically attempted to be preserved around intraoperatively identified language-positive sites. The grid was created in the tumoral and peritumoral areas. The pictures acquired from baselining were presented time locked to the TMS pulses while the coil was moved over the cerebral hemisphere. During this process, the patient was audiovisually recorded. A site was considered a positive “hit” for language eloquence if disruption (e.g., no response, performance error, and semantic error) was observed on any of the trials. A hesitation was counted when the patient’s response came after the TMS stimulation had ended for that picture. Positive sites were marked on the 3D brain surface as white dots. These sites were exported in digital imaging and communications in medicine (DICOM) files to be uploaded in the Neuronavigation system (Elements; BrainLab, Munich, Germany).

### Neuroimaging Acquisition & Preprocessing

Structural MRIs were performed using a 3T MRI scanner (GE Healthcare, Waukesha, Wisconsin) with an eight-channel head coil. DWI data was acquired using a diffusion-weighted spin-echo echo-planar imaging sequence (repetition time/echo time = 10 000/62 ms, matrix size = 128 × 128, field of view = 22 × 22 cm, slice thickness = 2.5 mm with no intersection gap, number of diffusion-weighting directions = 32, b value = 1000 s/mm2). In total, 44 slices were acquired, covering the medulla to the top of the brain. High-resolution 3D spoiled gradient-echo T1-weighted sequences were acquired for anatomic reference.

Preprocessing of DTI sequences was performed using Brainlab Elements (Brainlab AG; Munich, Germany). Brainlab Elements’ “Image Fusion” and “Distortion Correction” modules was also used to non-linearly co-register the preoperative T1 MRI used for navigated TMS mapping and intraoperative navigation with the postoperative T1 MRI (1 day postop). This elastic co-registration approach has been validated in multiple previous studies for correcting imaging distortions caused by surgically-induced brain shift.^21–24^

After preprocessing, white matter tract dissections were performed using a region of interest (ROI) approach. Each positive TMS point was defined as a ROI for fiber tracking in each patient. (Fig. 1D) The minimum fiber lengths of 30 mm and a maximum angulation of 50 degrees were used. Fiber tracts generated using TMS points as the seed are referred to as “TMS tracts” in this study. Given the significant inter-individual variability in fractional anisotropy (FA), a normalized approach for FA thresholding was used to standardize the selection to the individual profile of the patient.^25,26^ For each patient, the fractional anisotropic (FA) threshold was progressively increased by .01 after starting at 0 while holding all other parameters constant. When the threshold reached a value high enough to exclude all fibers, this value was defined as 100% for that patient. Four different percentages of FA thresholds were then analyzed across the whole cohort: 25%, 50%, 75%, 85%. Figure 1D illustrates TMS-based fiber tracking. Atlas-based segmentations from the Human Connectome Project (HCP842)^26^ were downloaded and used to obtain population-averaged volumes in MNI space. Both the binary masks of population-averaged tracts were obtained as well as the probabilistic templates that reflect the inter-individual variability of the cohort. TractSeg^27^ was used to produce data-driven fiber tract segmentations for the case example in Figure 7.

Cortical data (TMS points and associated connectivity-based subgroups of TMS points) and subcortical data (fiber tracts-“TMS tracts”) were segmented in Brainlab and exported. To facilitate accurate transformations in a cohort of brain tumor patients, a non-linear algorithm from SPM was used for spatial normalization to MNI.^11^ Matlab was used for voxel-wise summations to produce composite probabilistic atlases of TMS tracts and cortical points after normalization to MNI. In patients who had TMS tracts resected, Brainlab Elements was used to automatically segment the overlap between the tracts and resection cavity to produce tract-cavity segmentations. These segmentations were then normalized to MNI space to identify an approach for classifying “true positive” (resection of tract with corresponding deficit) and “false positive” (resection of tract without corresponding deficit) tract-cavity segmentations. A grid-search with k-fold cross-validation was used to define the optimal set of cutoffs for classification across three different measures of spatial association between tract-cavity segmentations and normative tract volumes.

The following cut-offs for measures of spatial association between resected TMS tract segments and normative tracts were analyzed within the grid search: Volume of overlap (cm^3): [.5, 1, 1.5, 2, 2.5] Number of normative tracts overlapped: [2, 3, and 4] and Percentage of overlap (% of tract-cavity segmentation overlapped by normative tract volumes): [40, 50, 60, 70, and 80]. The advanced normalization technique (ANTs) software package^28^ was used to warp normative data to patient space to apply these cutoffs to individualized predictions for the final iteration of the predictive model. The normalization transformations to MNI space were first computed. The inverse transformation was then used to warp HCP tracts from MNI to the patient space. The cutoffs for the three measures derived in MNI space were used to generate predictions within the final iteration of the predictive model reported in Figure 7.

For the analysis in Figure 8, the regions of co-localized TMS tracts and normative HCP tracts (“TMS-HCP”) that corresponded with a postoperative deficit (n=8) were segmented in MNI space to produce 8 regions-of-interest (ROIs). Each patient-specific ROI was analyzed for its degree of overlap with the normalized resection cavities from the whole cohort. If the spatial association between the patient-specific resection cavity and ROI exceeded the cutoffs defined in Figure 5, this was counted as a resection of that ROI for that patient. The same analysis was then performed along with the patient-specific TMS tract volumes. A resection was recorded if the cavity-ROI overlap included the TMS tract volume for that patient. The percentage of patients who had positive ROI resections that included TMS tracts versus patients who had positive ROI resections irrespective of TMS tracts were analyzed according to the percent who sustained long term deficits in each group.

### Resection Analyses

The resection cavity was segmented in a data-driven manner using the semi-automated “Smart Brush” from Brainlab Elements.^29^ Brainlab Elements’ “Image Fusion” and “Distortion Correction” modules were used to non-linearly co-register the preoperative T1 MRI used for navigated TMS mapping and intraoperative navigation with the postoperative T1 MRI (1 day postop). This elastic co-registration approach has been validated in multiple previous studies for correcting imaging distortions caused by surgically-induced brain shift.^21–24^ Overlap between preoperative reconstructions (TMS points, fiber tracts) and the resection cavity was automatically segmented using the “Intersection” function of the “Object Manipulation” module in Brainlab. For Figures 3, 4, and 5, overlap (versus no overlap) between the TMS points/tracts and the resection cavity were included as a binary predictor in binary logistic regression models that predict pre- to postoperative long-term language decline. For Figure 7 and Figure 8, the classification of resected tracts vs preserved tracts was based on the cutoff parameters derived in Figure 6 and shown below in the *Normalized Analyses* section of the Results.

### Statistical Analysis

Statistical analyses were performed using Python (Version 3.12.1). Scipy.stats, statsmodels, and pandas were used for basic computations such as t-tests and regression. Independent, one-tailed t-tests were utilized. Wald tests were used to evaluate the statistical significance of regression models. Likelihood ratio test (LRT) was used to evaluate the statistical significance of multivariate nested models compared to corresponding univariate models.

Seaborn and matplotlib were used for more complex visualizations such as scatter plots, box plots, and heatmaps. Scikit-learn generated ROC curves, confusion matrices, and k-fold cross-validation. Confusion matrix elements were used to analyze the sensitivity, specificity, positive predictive value (PPV), and negative predictive value (NPV) of the predictive models based on mapping and the resection volume. A true positive was defined as one or more resected sites (TMS points and/or white matter tracts (WMTs)) with a permanent deficit. A false positive was defined as one or more resected sites that did not have a corresponding permanent deficit. A true negative resulted from no sites being resected and no permanent deficit. A false negative was defined as having no sites resected but incurring a deficit.

## Results

Table 1 shows patient demographics, clinical characteristics, and surgical outcomes. Around 90% of patients had lesions defined as eloquent or near-eloquent according to Sawaya et al.^30^ Around a third of patients (27%) presented with aphasia (WAB AQ < 93.8) at baseline. About half (47%) experienced new or worsened aphasic deficits in the immediate postoperative period, and 10 patients (13%) sustained PLDs. Of the 10 patients with PLDs, three (30%) primarily had receptive deficits, while seven (70%) primarily had expressive deficits. With regards to the primary domain of neurocognitive impairment, three patients exhibited deficits in verbal fluency, one patient specifically had a deficit in semantic fluency, two had deficits in lexical retrieval (anomia), and two had deficits in auditory comprehension. Baseline demographics and clinical characteristics (whether binary or categorical- as shown in Table 1) were not significantly associated with PLDs using regression analysis. The group-level distribution of tumor volumes in MNI space demonstrated that tumors most commonly localized to the anterior and middle temporal lobe (Fig. 2).

**Figure 2:**
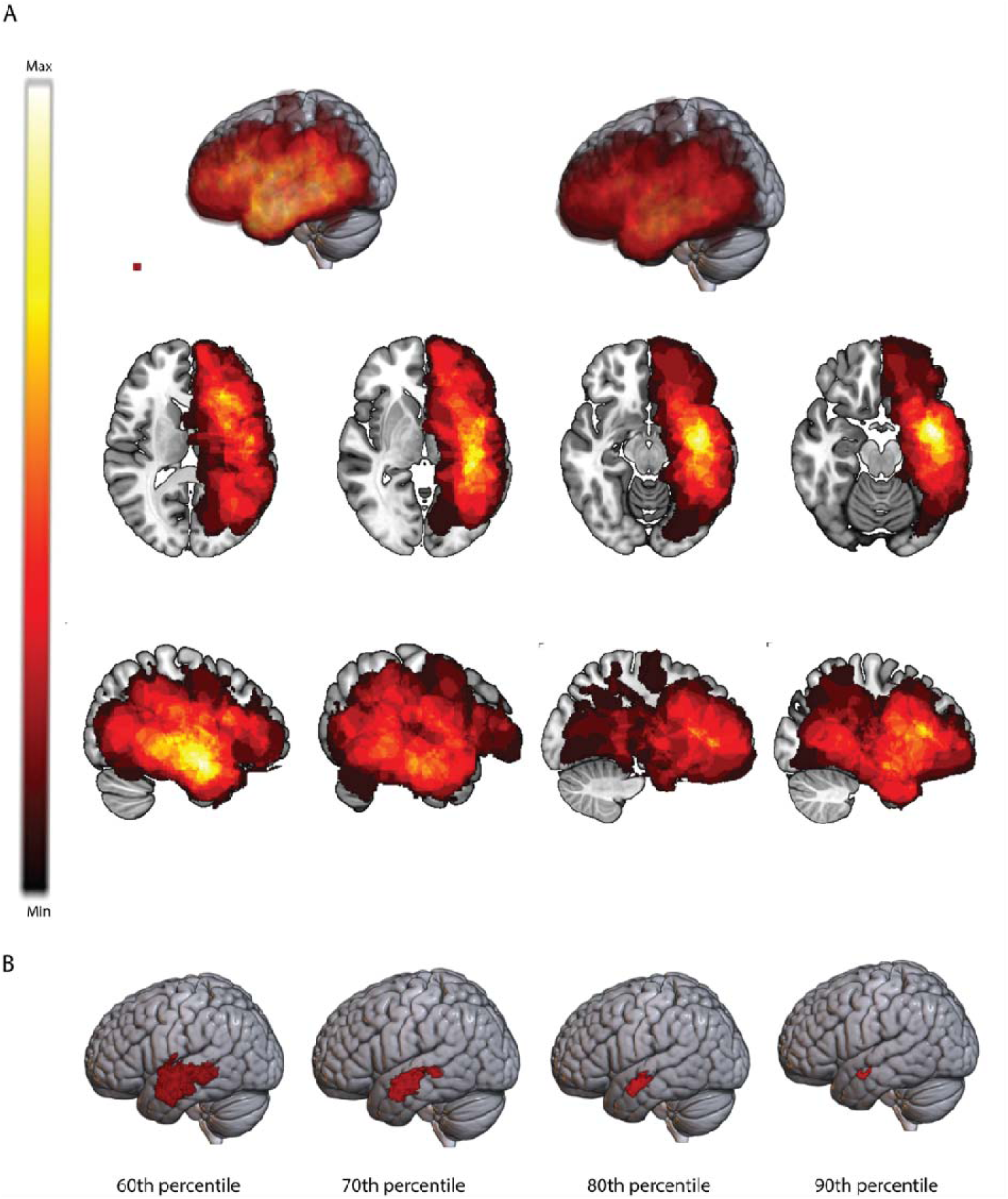
Probabilistic distribution of tumor volumes across the cohort of patients. (A) shows 3D rendering (top row) as well as axial (middle) and sagittal slices (bottom row). (B) shows the effects of progressive thresholding, showing that tumors most frequently localized to the anterior temporal lobe in this cohort.

**Figure 3:**
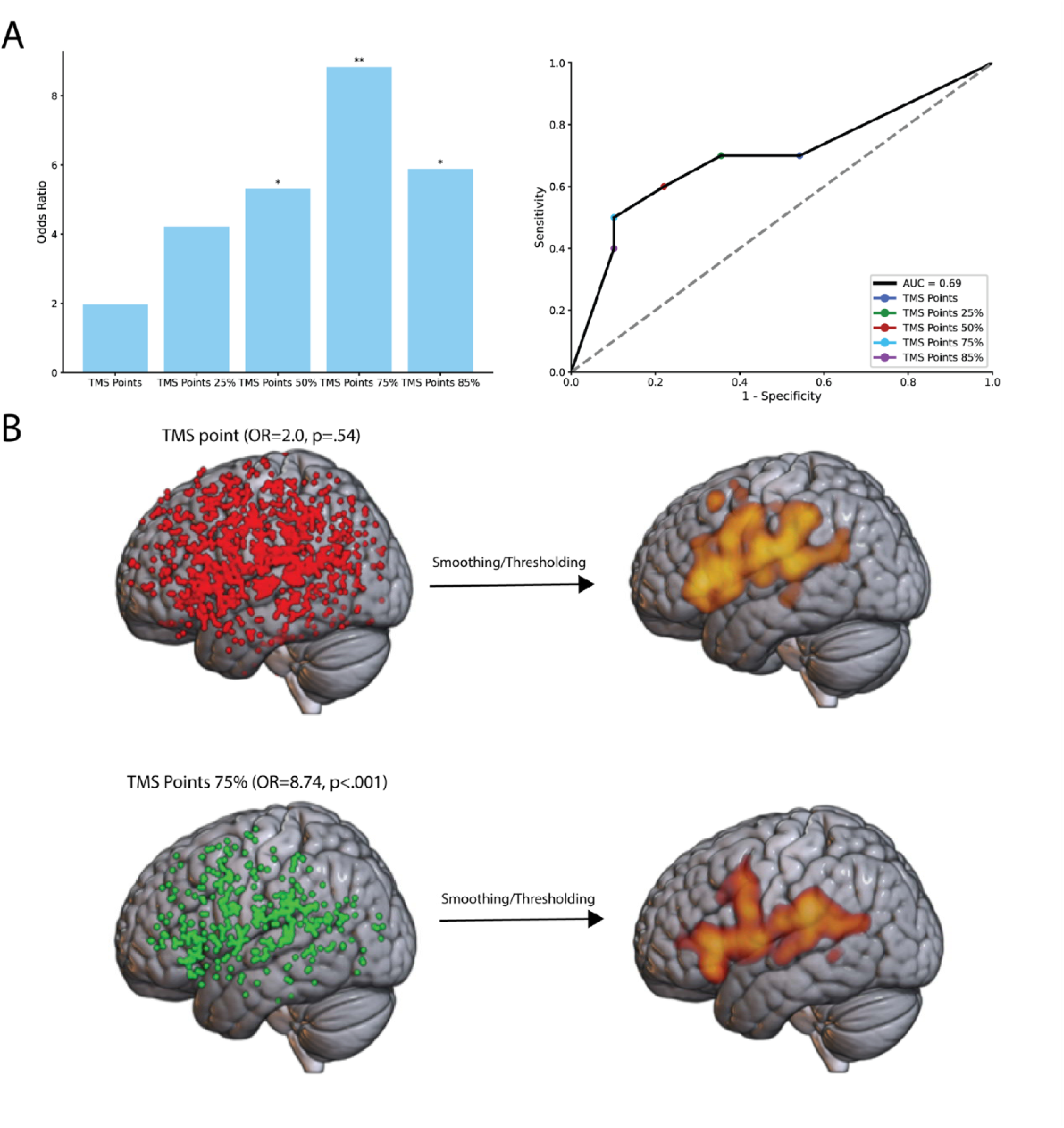
White matter connectivity determines the functionality of TMS data and defines a canonical distribution of cortical language sites at a group-level. (A) left shows the odds ratios for resecting different subgroups of TMS points defined by their white matter connectivity profile. (A) right shows the predictive value for resecting these subgroups with an ROC curve. (B) shows the cortical distribution of TMS points (left) and TMS points with the most predictive FA threshold of white matter connectivity (right). (D) shows the result of Gaussian smoothing (FWHM=10) and thresholding of cortical distributions of TMS points alone (left) and TMS points with robust white matter connectivity (right). (n=79. Wald test: *P < 0.05, <P < 0.01)

### Predicting Aphasia with Cortical Data

Resection of positive TMS points did not predict long-term aphasic deficits (OR=2.0, p=.54; Fig. 3A). Using TMS points as the cortical ROI for fiber tracking, we created subgroups of TMS points according to the fractional anisotropy of the tracts connected to each point. For example, points with connecting tracts at the 75% FA threshold (“TMS points 75%”) were grouped differently than TMS points with tracts connecting at lower FA thresholds. PLDs significantly associated with the resection of TMS point subgroups of white matter connectivity at FA thresholds of 50% and above, with the greatest odds ratio found for the TMS points 75% group (OR=8.74 , p=<.01; Fig. 3A- left). An ROC curve for predicting PLDs with TMS points using FA as the classification threshold demonstrated an Area Under the Curve (AUC) of .69 (Fig. 3A- right). The most predictive TMS point subgroup (TMS points 75%) demonstrated a group-level cortical distribution that closely aligns with canonical patterns derived from invasive stimulation: hotspots in the frontal operculum, ventral precentral gyrus, and posterior superior temporal gyrus (Fig. 3B).^31^

### Predicting Aphasia with Subcortical Data

We then analyzed the WMTs produced by TMS-based fiber tracking by analyzing their predictive value for PLDs at different FA thresholds. Resection of the tracts reconstructed at the 75% FA threshold were the most predictive (OR=60.0, p<.001; Fig. 4A- left). Figure 4A shows an ROC curve for TMS tracts in the same format as the cortical ROC curve in Figure 3, demonstrating a much more robust Area Under the Curve (AUC=.90; Fig. 4A- right). We show that these tracts recapitulate normative patterns of white matter connectivity at a group-level by normalizing them to MNI space and summing together across patients to create a probabilistic anatomical distribution. This probabilistic atlas showed subcortical hotspots (high voxel values indicating that many patients had a tract localize to that voxel) that anatomically co-localize with normative, language associated WMTs defined by the Human Connectome Project^26^ (HCP) (Fig. 3A). The highest value voxels of both the TMS and HCP atlases localized to the posterior segment of the arcuate fasciculus (Fig. 4C). Additionally, the group-level TMS tracts in MNI space reproduce the same voxel value pattern of the SLF as probabilistic group-level mapping of the SLF in healthy subjects from the HCP (Fig. 4D). Both have significantly lower voxel values anterior to the precentral gyrus, demonstrating significantly more inter-individual variability in the frontal region.

**Figure 4:**
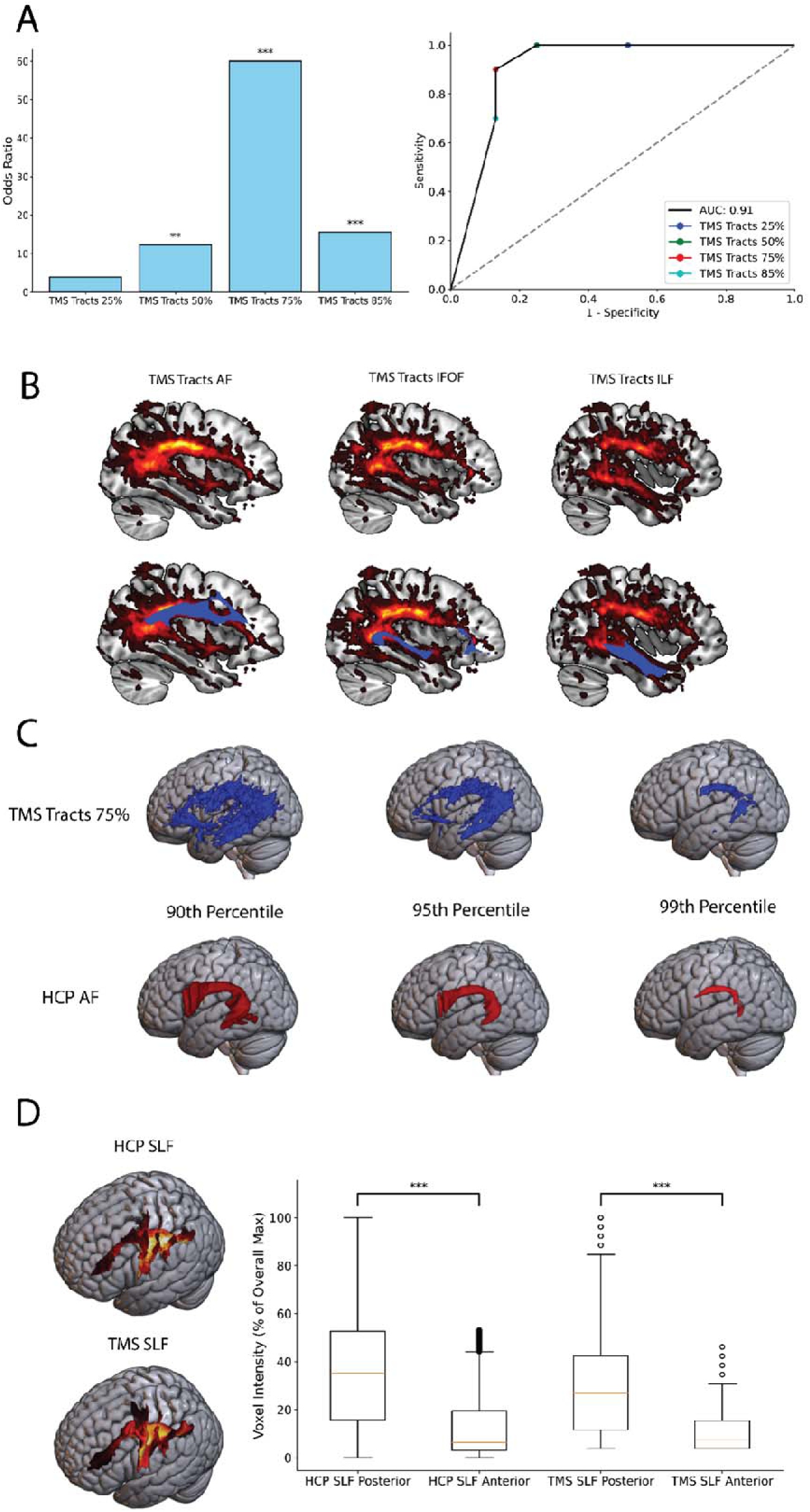
D**a**ta**-driven fiber tracking based on cortical language mapping in glioma patients localizes patient-specific WMTs that predict PLDs and recapitulate healthy normative patterns of white matter connectivity at a group-level.** (A) shows the odds ratios for resecting TMS tracts according to the FA threshold used for their reconstruction (left) while the right figure shows the predictive value for resecting these tracts at different FA thresholds with an ROC curve. (B) shows the probabilistic spatial distribution of TMS tracts normalized to MNI space and summed with a voxel-wise summation. Top row shows the overlay of language- associated WMTs in MNI space from the HCP project, depicting how TMS tract hotspots co- localize with normative tract segmentations. (C) shows progressive thresholding for probabilistic TMS tracts from this study compared to progressive thresholding for the probabilistic arcuate fasciculus from the HCP project. Both converge along the posterior parieto-temporal segment of the arcuate fasciculus. (D) Shows the comparison between the probabilistic SLF from the HCP project and the SLF portion of the probabilistic TMS tracts. The visualization shows that hotspots localize to the posterior segment of the tract in both cases. The box plot shows the average voxel intensities for the anterior (anterior to the precentral gryus) and posterior (precentral gyrus and posterior) segments of the SLF in both the HCP and TMS probabilistic reconstructions. (n=79. One-tailed t-tests, linear regression: \**P* < 0.05, <*P* < 0.01, <\**P* < 0.001. Error bars represent SD)

**Figure 5:**
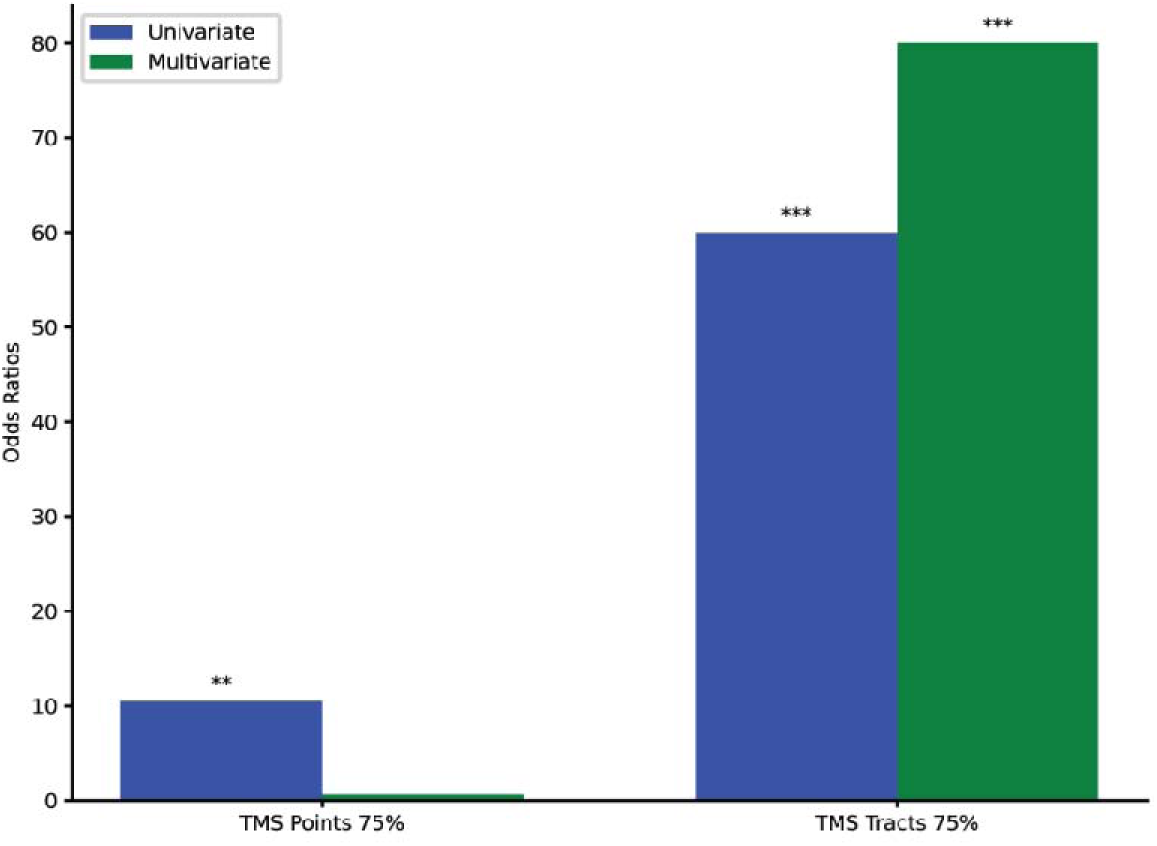
Subcortical damage to TMS tracts independently predicts PLDs. This figure depicts univariate versus multivariate analysis for prediction of PLDs. Multivariate analysis included both subcortical tracts and the most predictive subgroup of TMS points. This plot shows that adding cortical data to the subcortical model does not improve the predictive value in a statistically significant manner while adding subcortical data to the cortical model does significantly decrease its predictive value. (\**P* < 0.05, <*P* < 0.01, <\**P* < 0.001)

**Figure 6:**
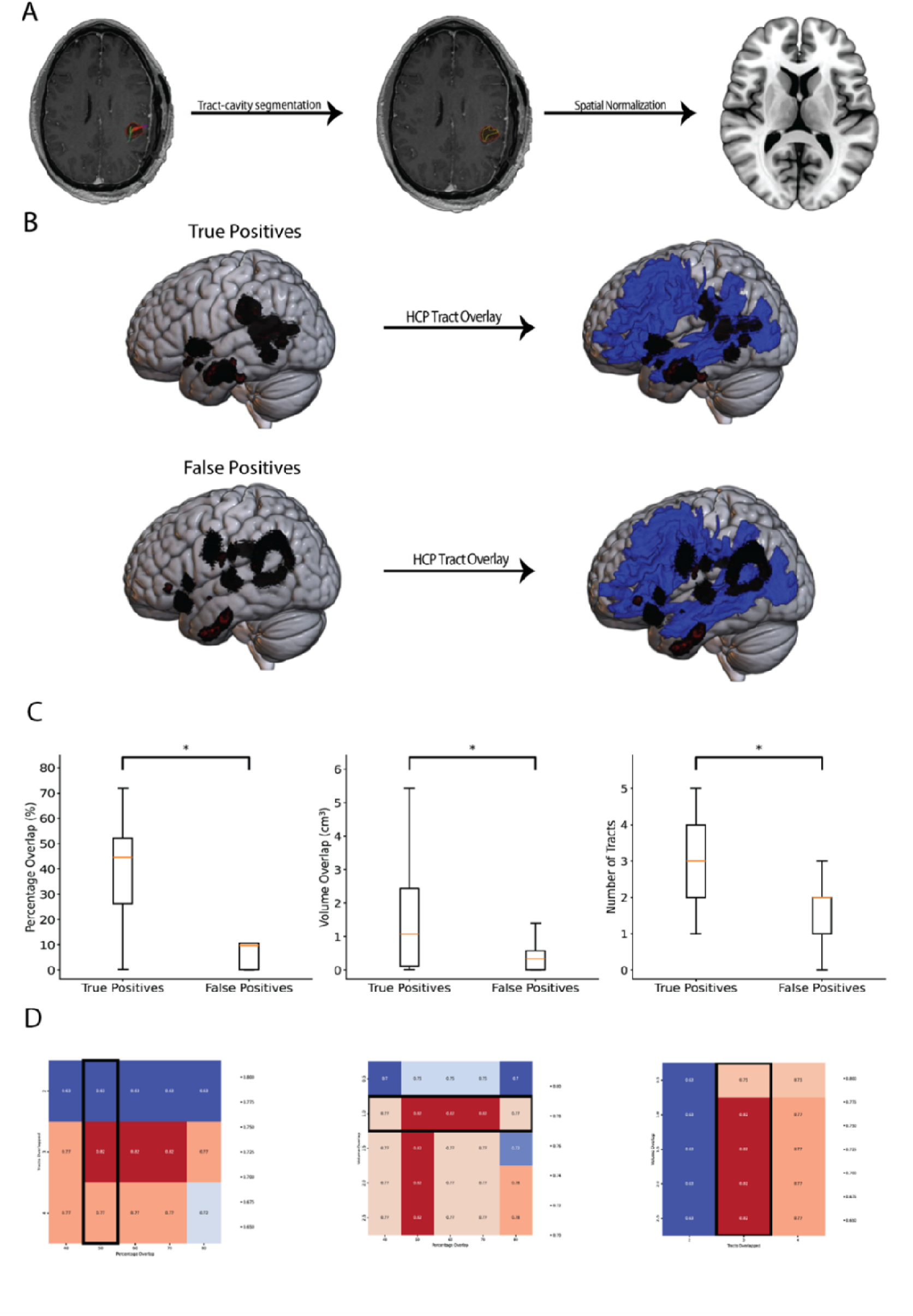
N**o**rmalized **analysis. The resected segment of TMS tracts in patients who sustained PLDs (true positives) co-localize with normative HCP language-associated tracts in MNI space compared to resected tract segments in patients without PLDs (false positives).** (A) shows the tract-cavity segmentation and spatial normalization workflow (top left) along with a composite visualization of the true positive and false positive tract-cavity segmentations in MNI space (top right and bottom). (B) shows a visualization of the spatial association between resected tract segments and normative tracts. (C) shows that true positives have significantly more co-localization with normative, language-associated tracts from the HCP than false positives. (D) shows heatmaps that illustrate the grid-search process to find the optimal cutoffs across the three measures that most accurately classify true positives versus false positives. The black rectangle indicates the row or column that showed the highest accuracy for that heatmap. This process resulted in the selection of these three parameters: over 1 cm^3 volume of infiltration, more than 3 tracts overlapped, and more than 50% of the tract-cavity segmentation overlapped. (n=18. One tailed t-test: \**P* < 0.05)

**Figure 7:**
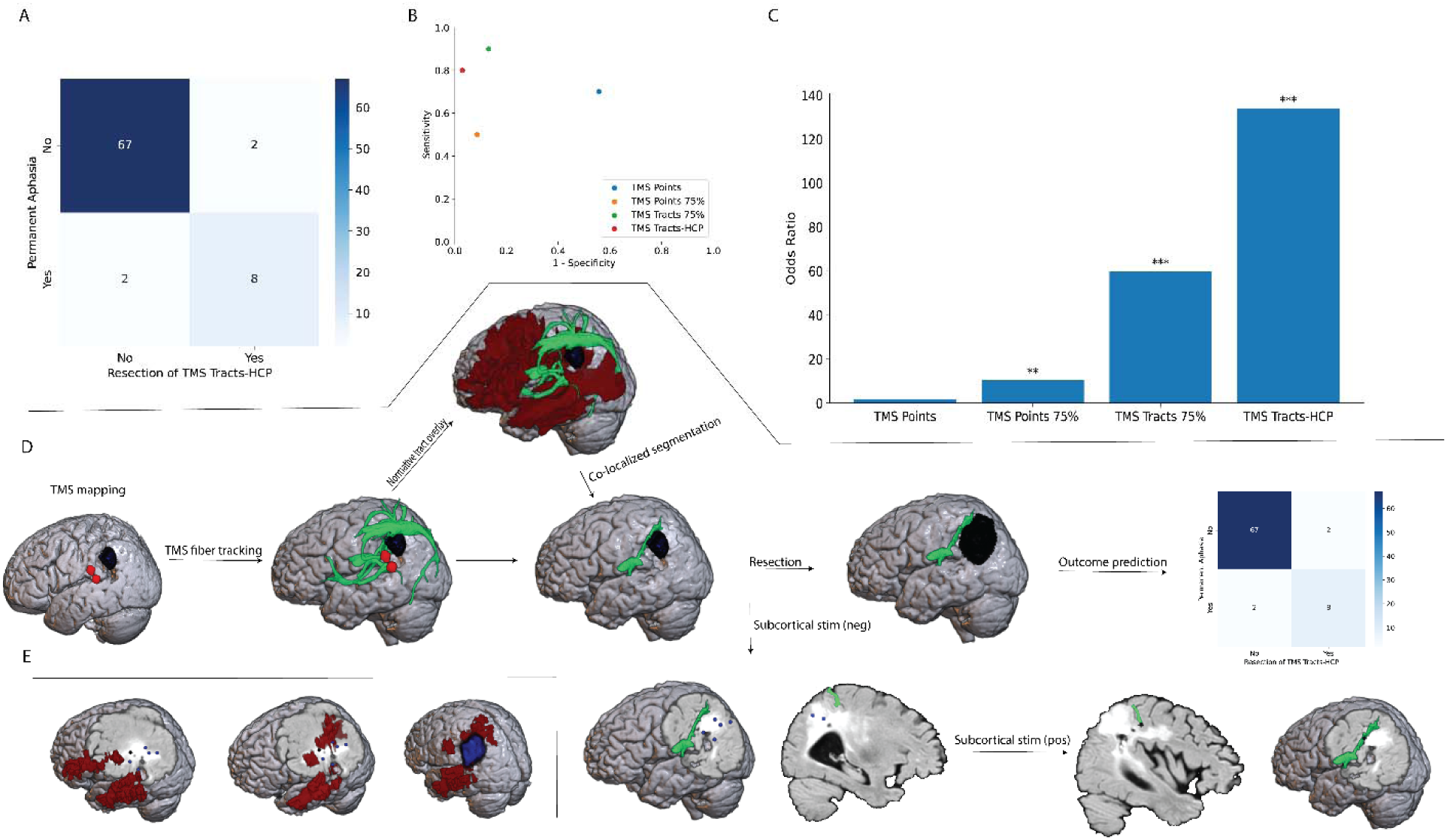
O**v**erview **of the development process, final performance of predictive model, and prospective validation with direct electrophysiological stimulation.** (A) shows a confusion matrix for the best performing final model in predicting PLDs. (n=79). (B) shows how the odds ratio significantly increases with each iteration, culminating in an odds ratio of 134 in the final comprehensive model. (C) shows the predictive value of each of the developing components of the model in terms of an ROC plot. (D) illustrates the prospective application of the data for surgical planning and execution with a flow chart. Patient presents with a dominant hemisphere lesion and undergoes TMS mapping and fiber tracking. Normative tracts are warped to patient- space and overlayed onto individualized TMS tracts to produce a segmentation of co-localized voxels. The co-localized segment accurately predicted positive and negative subcortical stimulation responses (n=6), extent of resection (n=1), and surgical outcome (n=1). (E) shows the poor performance of TractSeg white matter segmentations in predicting the same metrics. Left shows the AF (red) with the negative (blue) and positive (black) stimulation point. Middle shows the MdLF (red) with the negative and positive stimulation points. Right shows the significant overlap between the resection cavity (blue) and MdLF. (Wald test: \**P* < 0.05, <*P* < 0.01, <\**P* < 0.001)

**Figure 8:**
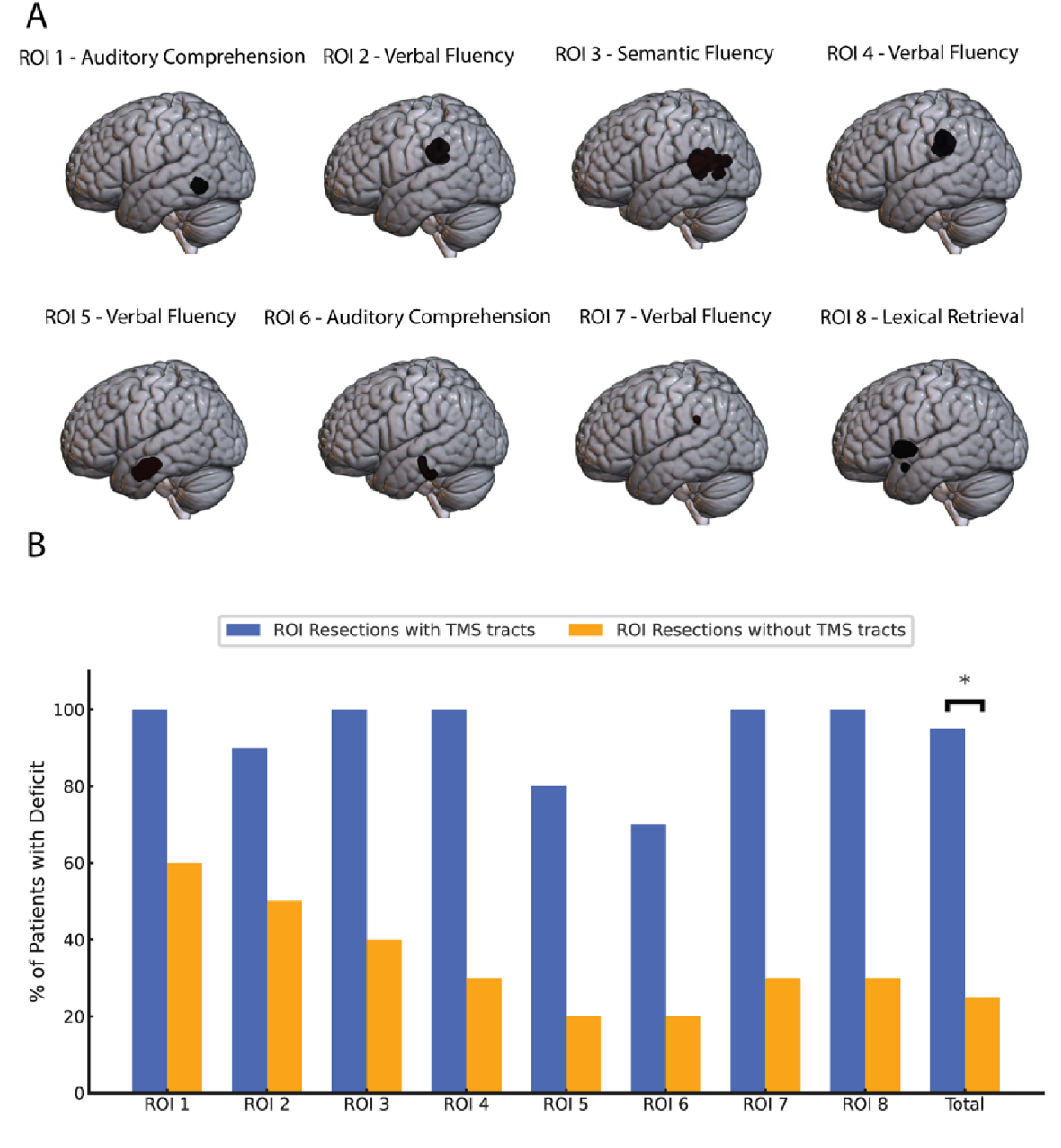
Normalized analysis. PLDs result from surgical damage to *individualized* ROIs within language-associated WMTs. (A) Resecting the anatomical region defined by these ROIs only compromises language function if co-localized TMS tracts are resected. Damage to the anatomical region defined by these ROIs does not predict outcomes at a group-level without individualized data. (B) Patient-specific anatomical locations of the resected co-localized TMS tract-HCP segmentations (smoothed, FWHM=5) in patients with PLDs. (n=8. One-tailed t-test: *P < 0.05)

Multivariate analysis (Fig. 5) with the most predictive cortical sites (TMS points 75%; OR=8.74, p=.0021) and subcortical tracts (TMS tracts 75%; OR=60.00, p<.001) showed that the resection of TMS points 75% no longer predicted PLDs (OR=.62, p=.63). However, the resection of TMS tracts remained a significant predictor (OR=80.00, p<.001) in multivariate analysis, indicating that TMS tracts predict PLDs independently of the most predictive subgroup of cortical language sites. The Likelihood Ratio Test (LRT) demonstrated that the addition of cortical data to the subcortical model did not significantly decrease its predictive value (χ=.23, p=.63) but the addition of subcortical data to the cortical model significantly decreased its predictive value (χ=16, p<.001). Surgical damage to TMS tracts reconstructed at the most predictive FA threshold demonstrated a sensitivity of 90%, specificity of 87%, PPV of 50%, and an NPV of 98% for predicting PLDs. This analysis indicates that aphasic deficits result from surgical damage to TMS-derived fiber tracts, though the cortical sites themselves do not seem to contribute to functional decline.

### Normalized Analyses

To address the significant number of false positives (TMS tract resection with no deficit), we normalized the resected portion of TMS tracts (“tract-cavity segmentations”) in each patient to MNI (Fig. 6A). Initial composite visualization of true positives (n=9) and false positives (n=9) in MNI space revealed that true positives cluster within the white matter of the anterior temporal lobe and posterior temporoparietal junction (TPJ) (Fig. 5B). In contrast, false positive tract segments distributed throughout the left hemisphere and did not show a discernable pattern.

In contrast to false positive segmentations, true positive tract-cavity segmentations co- localized with the normative volumes of HCP-defined, language-associated WMTs (Fig. 6B). True positive segmentations demonstrated significantly higher spatial association with normative tracts compared to false positives across all three measures: volume of overlap (p<.05), percentage of overlap (p<.05), and number of normative tracts overlapped (p<.05). Additionally, every patient had at least one atlas-based tract segmentation resected (not shown), demonstrating the inadequacy of normative data and the importance of individualized mapping. This analysis shows that essential function localizes to subregions within normative language-associated WMTs.

We hypothesized that measures of normative tract association (overlap volume, percentage, and number of tracts) could classify resections of TMS tracts to improve the predictive model for PLDs. We computed the classification performance of every possible permutation of cutoffs for classifying TMS tract resections with these measures (Fig. 6D). We computed and highlighted the row or column of each heatmap that showed the highest average accuracy, indicating the best cutoff for each respective measure. This process resulted in selecting the following three parameters as the best predictors of PLDs: over 1 cm^3 volume of infiltration, more than 3 tracts overlapped, and more than 50% of the tract-cavity segmentation overlapping. These cutoffs were then later applied to make predictions on an individual level.

### Final Predictive Model

To perform the analysis for the final iteration of the predictive model, we warped the normative volumes of the 7 language-associated tracts (AF, SLF, FAT, IFOF, ILF, MdLF, UF) from MNI space to patient-specific anatomy (n=79). The resection volume, normative HCP tracts, and individualized TMS tracts (and derived cutoffs) were used to generate outcome predictions in each patient. This resulted in 80% sensitivity, 97% specificity, 80% PPV, and 97% NPV (Fig. 7A) for predicting PLDs. The odds ratio significantly increased with each iteration, culminating in an odds ratio of 134 (CI: 16-1086, p<.001) (Fig. 7B).

Figure 7D illustrates the workflow for the prospective application of the predictive model along with its validation with DES. The patient presents with a lesion in the dominant hemisphere and undergoes navigated TMS language mapping and fiber tracking with notable spurious streamlines. Normative tracts are warped to the patient’s anatomy and overlayed to provide data-driven filtering of TMS tracts. The complete preservation of co-localized segments correlated with intact postoperative language function, demonstrating accurate outcome prediction for this patient.

This flow chart also shows that the co-localized tract segmentation predicted this patient’s real-time linguistic performance (error vs task completion) in response to direct electrophysiological stimulation of the subcortical brain. As the resection progressed, the surgical cavity did not elicit errors until the co-localized segmentation was reached. Here, subcortical stimulation at 5 milliamps elicited strikingly reproducible semantic errors associated with an object naming paradigm. The positive stimulation point co-localized with the co-localized segmentation at a distance of 0 millimeters. The negative stimulation points (n=6) were more than 5 mm from the co-localized segmentation.

In contrast, state-of-the-art automated fiber tracking (TractSeg) did not accurately predict intraoperative mapping or the extent of resection/postoperative outcome (Fig. 7E). The left image shows how the positive stimulation point localizes distantly from the AF segmentation (in contrast to the TMS-HCP co-localized segmentation). Although the positive stimulation point did co-localize with the middle longitudinal fasciculus (MdLF, anterior temporal - superior temporal - parietal conenction), two negative points also co-localized with this tract. Additionally, a significant portion of the MdLF was resected, indicating an inaccurate outcome prediction based on TractSeg for this patient.

### Importance of Individualized Mapping

To investigate whether PLDs can be independently predicted by normative anatomy, we created anatomical ROIs in MNI space from the co-localized segments of individualized and normative tracts in patients whose resections accurately predicted aphasic decline (n=8). Figure 8A shows the anatomical location of each ROI along with the associated neurocognitive deficit. Using normalized resection cavities (n=79), we analyzed the group-level association of these regions with permanent deficits (if resected) with and without co-localized TMS tracts. 92% of patients who underwent resections that included TMS tracts sustained deficits, while 34% of patients who underwent ROI resections, irrespective of individualized data, sustained deficits (p<.05) (Fig. 8B).

## Discussion

To our knowledge, these results constitute the first prediction of linguistic outcomes based on data-driven imaging and individualized resection volumes. Our approach’s accuracy (>90%) and personalized standardization confer unique clinical value. This approach identifies the cortical regions and white matter pathways specific to each patient that are necessary to sustain language function. This data could not only decrease long-term deficits but also increase rates of gross total resections.^32^ Instead of settling for conservative approaches due to uncertainty, surgeons could gain data-driven confidence to safely extend resections to improve survival. Using this non-invasive mapping technique, surgeons could preoperatively “simulate” the functional outcome of surgical approaches to select a personalized strategy to achieve maximal safe resection. The multimodal method could also optimize non-invasive treatments for glioma patients, such as radiation therapy for tumor control or brain stimulation for language rehabilitation.

This study provides a novel anatomical explanation for the origins of aphasic surgical deficits. Here we show that essential language function localizes to the subcortical level, specifically within individualized segments of language-associated WMTs. Figure 8 demonstrates that these white matter subregions require individualized mapping for surgical preservation, consistent with the confusing anatomical heterogeneity of postoperative aphasia reported by neurosurgeons.^33,34^ The resected portion of tracts in patients who sustained PLDs clustered into dual anterior/posterior anatomical foci of aphasic vulnerability: one deep to the anterior temporal lobe (ATL) and another deep to the temporoparietal junction (TPJ). These regions have a high density of white matter pathways that serve the language system and its embedding with other cognitive functions.^35^ This finding remarkably recapitulates results reported in the stroke and resection lesion mapping literature that also identify these two regions as strongly correlated with PLDs.^36–38^

This study demonstrates that white matter connectivity plays a significant role in defining the cortical language network’s function and functional anatomy in glioma patients. While TMS points alone showed no functional relevance, specific subgroups defined by white matter connectivity demonstrated consistent functionality. In contrast to unfiltered TMS points, the most predictive of these connectivity subgroups clearly demonstrated group-level clusters in the frontal operculum, ventral precentral gyrus, and posterior superior temporal gyrus. Notably, this non-invasively captured pattern closely recapitulates the cortical distribution of DES-positive language sites reported in the most extensive study of intraoperative language mapping to date.^31^

Figure 5 not only shows that subcortical tract injury independently predicts deficits but also demonstrates that cortical resection data does not improve the predictive value of the subcortical model. These results fit well with previous studies. Duffau et al have repeatedly written about the importance of preserving white matter connectivity to maintain capacity for long term function.^39,40^ Additionally, a landmark study in the stroke literature found that cognitive deficits primarily result from subcortical disconnection rather than cortical damage, while recent stroke studies have predicted a wide array of cognitive-behavioral deficits based on white matter disconnections.^37,41^ Studies have shown that while cortical topology has significant potential for neuroplastic reorganization and functional compensation, subcortical circuitry seems to lack this dynamic flexibility.^39,40,42^

Prospective application of the individualized and normative data together precisely predicted the results of direct electrophysiological stimulation of subcortical brain in a patient with recurrent glioblastoma. The accurate, patient-specific representation of essential function within severely distorted anatomy illustrates how the synergy of this imaging approach allows for strong outcome predictions in an anatomically diverse cohort of surgical glioma patients. When warped to patient-space, the large normative tract volumes supply a patient-specific foundational architecture of connectivity to constrain the individualized TMS tracts that often include spurious fibers. While this normative architecture significantly overrepresents relevant white matter, individualized data used in tandem can map its patient-specific functional subregions. TractSeg, a sophisticated deep learning approach to data-driven fiber tracking, demonstrated clear deficiencies in localizing intraoperatively identified subcortical sites of functionality, underscoring the necessity of integrating clinically relevant outcome measures into the brain mapping development process.

This study, along with previous work from our group on predicting motor outcomes, reveals the clinical utility of using fractional anisotropy (FA) as an imaging biomarker for long-term functionality.^43–45^ Figure 4 illustrates this by showing a robust ROC curve using DTI to predict PLDs with FA as the classification threshold. This likely reflects activity dependent myelination coupled with dynamic changes in neural activity induced by the tumor. Increased activity in certain tract segments leads to increased myelination and corresponding increases in FA.^46^ FA-based thresholding approaches to tractography may selectively localize the unique tract segments that each patient depends on to sustain language processing given the specific constraints imposed by the tumor.

While tumor invasion and associated edema can decrease fractional anisotropy, these insults can often render the tract non-functional as well, supporting the use of FA as a biomarker.^47,48^ However, it is important to note that the predictive value of this metric strongly depends on normalization to the individual anisotropic profile of each patient, which transforms the absolute values commonly used in the literature to the percentages outlined in this study.^25^ Additionally, the approach to cortical seeding significantly impacts the predictive value of FA, as demonstrated in our previous work that shows a much weaker AUC for predicting motor outcomes using anatomical seeds versus functional. Future work should fully characterize the role of FA as an imaging biomarker for long-term function as well as seek to identify synergistic markers.

FA correlates positively with streamline directionality and inversely with fiber complexity, meaning that regions of higher FA generally reflect more uniform diffusion and fewer crossing fibers.^49,50^ This characteristic may explain why our DTI-based models achieve strong predictive accuracy for PLDs despite their inability to fully resolve complex fiber orientations. Although crossing fibers are anatomically widespread, they may not independently contribute to PLDs. Indeed, a previous stroke-based study using DTI models identified the same vulnerable regions in the anterior temporal lobe (ATL) and temporoparietal junction (TPJ) that we report here, despite similar technical limitations. ^38^ These findings may suggest that complex fiber configurations provide structural redundancy by distributing linguistic processing across multiple local pathways. In contrast, uniformly oriented fiber segments with high FA values may represent “bottlenecks” for essential, non-redundant linguistic information transmission, rendering them more functionally susceptible to surgical damage. Furthermore, our study shows that postprocess filtering with normative data can effectively compensate for false positive reconstructions resulting from the limitations of patient-specific DTI data. Together, these results highlight the potential for DTI-based approaches to accurately predict PLDs. Future research should further investigate the role of crossing fibers in sustaining language processing and contributing to PLDs.

The cutoff parameters derived in Figure 6D likely do not reflect biological reality but rather serve as a proof-of-concept that the association between preoperative imaging and the resection volume can predict the functional impact of surgical strategies. We speculate that these cutoffs are effective because they accommodate for slight errors in the image processing pipeline, maximizing the statistical probability that positive predictions by the model accurately correspond to meaningful resection of the region. Similarly, we believe that the atlas-based tract segmentations are effective because they provide statistical filters for unconstrained individualized mapping data, not because their volumes accurately represent patient-specific white matter connectivity. Despite possible concerns regarding the accuracy of spatial normalizations in brain tumor patients, these data show that non-linear transformations perform well enough to allow normative data to classify the functional and non-functional segments of individualized TMS tracts in glioma patients. This finding supports previous studies showing the efficacy of various techniques for spatial normalization in patients with brain lesions.^11,42,51^

The neurosurgical literature has extensively described the phenomenon of “brain shift”- intraoperative mechanical deformation that can distort image-guided navigation.^52,53^ Some assert that this confounds the position of preoperative localizations relative to the postoperative resection volume, obfuscating which structures were removed. However, the conclusions of our study rely specifically on the consistent perioperative localization of WMTs, which reside in subcortical regions much less susceptible to deformation than the cortical surface.^54–56^ Additionally, multiple recent studies have shown that elastic, non-linear co-registration algorithms efficaciously correct for intraoperative deformation.^54,57,58^ These data, combined with the robust predictive models reported in this study, indicate that computational techniques can sufficiently address any perioperative anatomical distortions in glioma patients that may confound long-term outcome predictions based on individualized resection volumes.

Previous work can draw limited conclusions regarding the structure of functional subcortical networks. While many established neuroimaging techniques can interrogate global cortical function, insights into subcortical network underpinnings derive from either functionally ambiguous DTI tractography or focal direct stimulation limited by the surgical window. This limits the ability of researchers to map entire segments of functional WMTs, obfuscating the 3D spatial organization of functional subcortical fascicles.^59^ We address these technical neuroscientific constraints by applying a double layer of functional integration to white matter tract reconstruction from diffusion-weighted imaging.

First, cortical functional sites from focal brain stimulation were used as seeds for fiber tracking. Next, we functionally optimized these reconstructions by finding the postprocessing pipeline that maximizes the association between network damage and PLDs across a large cohort. This pipeline includes the individualized FA threshold selection approach along with data-driven streamline filtering using normative HCP data. This iterative process culminated in the development of a novel, data-driven brain mapping approach capable of non-invasively mapping the functionally essential components of subcortical networks. In addition to being a robust clinical tool for informing surgical decisions, the reproducibility, accessibility, and scalability of this technique make it a promising approach for researchers to investigate cognitive functional structure in the context of gliomas. The method significantly improves upon existing white matter reconstruction techniques by providing representations of entire segments of language-essential white matter without the hazards of invasive procedures.

While many studies have generated computational models to predict the outcome of epilepsy surgery and inform clinical decisions, this work represents a first-of-its-kind for glioma surgery.^60^ We show that preoperative TMS mapping, combined with diffusion imaging and normative data, can accurately predict the functional outcome of volumetric surgical strategies. Given that co-localizations between individualized and normative tracts can predict long-term language ability when combined with measures of surgical damage, this personalized data can reveal one of the most important and elusive^11,34^ characteristics for dominant hemisphere gliomas: proximity to critical language regions. The novel imaging method could significantly improve patient prognostication and stratification in clinical trials, contributing to the scaled development of new personalized treatment strategies for glioma patients.

### Limitations

This study has several areas for improvement. To facilitate rapid clinical application, we chose to use a widely available clinical neuronavigation software (Brainlab Elements) that uses a diffusion tensor imaging algorithm without the ability to resolve crossing fibers. Future work should utilize more flexible and advanced fiber tracking software to directly analyze the effect of higher resolution algorithms such as CSD in predicting PLDs.^61^ In addition, using the TMS- positive sites as ROIs to seed the WMT propagation limits the reconstruction of white matter to the functionally implicated streamlines in the peritumoral region rather than representing the full pathway. Retrospective clinical studies in neurosurgery face inherent limitations, including selection bias, incomplete data, and reliance on pre-existing records, which can compromise the validity of findings. Future studies should prioritize prospective designs with standardized data collection and well-defined patient selection criteria to minimize bias and strengthen the reliability and applicability of results. Finally, predictive models developed without independent validation may become overfit to the specific characteristics of the study population. Future work should utilize cohorts large enough to be divided into training and validation subsets to integrate measures of generalizability into the development process.

## Conclusion

This study demonstrates that long-term postoperative language deficits in glioma patients result from resecting individualized segments of critical white matter tracts. We integrate these findings into a personalized approach that uses cortical functional data and routine imaging to preoperatively predict the impact of individual tumor surgeries on language function with an accuracy of 94%. This study highlights the novel potential to preoperatively tailor surgical strategies using data-driven predictive modeling to achieve maximal safe resection.

### Author Contributions

MM: Study concept and design; acquisition, analysis, and interpretation of data; drafting and revising of the manuscript. KN: analysis and interpretation of data; revising manuscript. SP: data acquisition, administration, and revising manuscript. HM: acquisition of data. JT: revising manuscript. VK: revising manuscript. HL: revising manuscript. CE: revising manuscript. SF: acquisition of data. JW: acquisition of data. FL: acquisition of data. BT: revising manuscript. SS: Study concept and design, acquisition of data, revising manuscript.

### Data Availability

The institutional-specific raw data from this study is available upon reasonable request. The publicly available normative tract volumes can be accessed at: https://brain.labsolver.org/hcp_trk_atlas.html

## References

1. Schaff LR, Mellinghoff IK. Glioblastoma and Other Primary Brain Malignancies in Adults: A Review. Jama. 2023; 329(7):574–587.

2. Marko NF, Weil RJ, Schroeder JL, Lang FF, Suki D, Sawaya RE. Extent of resection of glioblastoma revisited: personalized survival modeling facilitates more accurate survival prediction and supports a maximum-safe-resection approach to surgery. J Clin Oncol. 2014; 32(8):774–782.

3. Ius T, Isola M, Budai R, et al. Low-grade glioma surgery in eloquent areas: volumetric analysis of extent of resection and its impact on overall survival. A single-institution experience in 190 patients: clinical article. J Neurosurg. 2012; 117(6):1039-1052.

4. Rahman M, Abbatematteo J, De Leo EK, et al. The effects of new or worsened postoperative neurological deficits on survival of patients with glioblastoma. J Neurosurg. 2017; 127(1):123–131.

5. Herholz K, Thiel A, Wienhard K, et al. Individual functional anatomy of verb generation. Neuroimage. 1996; 3(3 Pt 1):185-194.

6. Ojemann G, Ojemann J, Lettich E, Berger M. Cortical language localization in left, dominant hemisphere. An electrical stimulation mapping investigation in 117 patients. J Neurosurg. 1989; 71(3):316-326.

7. Ojemann GA, Whitaker HA. Language localization and variability. Brain Lang. 1978; 6(2):239–260.

8. Duffau H, Mandonnet E. The "onco-functional balance" in surgery for diffuse low-grade glioma: integrating the extent of resection with quality of life. Acta Neurochir (Wien). 2013; 155(6):951–957.

9. Price CJ. The anatomy of language: a review of 100 fMRI studies published in 2009. Ann N Y Acad Sci. 2010; 1191:62–88.

10. Giussani C, Roux FE, Ojemann J, Sganzerla EP, Pirillo D, Papagno C. Is preoperative functional magnetic resonance imaging reliable for language areas mapping in brain tumor surgery? Review of language functional magnetic resonance imaging and direct cortical stimulation correlation studies. Neurosurgery. 2010; 66(1):113–120.

11. Ius T, Angelini E, Thiebaut de Schotten M, Mandonnet E, Duffau H. Evidence for potentials and limitations of brain plasticity using an atlas of functional resectability of WHO grade II gliomas: towards a "minimal common brain". Neuroimage. 2011; 56(3):992–1000.

12. Sanai N, Mirzadeh Z, Berger MS. Functional outcome after language mapping for glioma resection. N Engl J Med. 2008; 358(1):18–27.

13. Mandonnet E, Winkler PA, Duffau H. Direct electrical stimulation as an input gate into brain functional networks: principles, advantages and limitations. Acta Neurochir (Wien). 2010; 152(2):185–193.

14. Muir M, Patel R, Traylor J, et al. Validation of Non-invasive Language Mapping Modalities for Eloquent Tumor Resection: A Pilot Study. Front Neurosci. 2022; 16:833073.

15. Ille S, Sollmann N, Hauck T, et al. Combined noninvasive language mapping by navigated transcranial magnetic stimulation and functional MRI and its comparison with direct cortical stimulation. J Neurosurg. 2015; 123(1):212–225.

16. Ruohonen J, Karhu J. Navigated transcranial magnetic stimulation. Neurophysiol Clin. 2010; 40(1):7–17.

17. Kertesz A. Western Aphasia Battery-Revised. The Psychological Corporation. 2007.

18. Lioumis P, Zhdanov A, Mäkelä N, et al. A novel approach for documenting naming errors induced by navigated transcranial magnetic stimulation. J Neurosci Methods. 2012; 204(2):349–354.

19. Krieg SM, Lioumis P, Mäkelä JP, et al. Protocol for motor and language mapping by navigated TMS in patients and healthy volunteers; workshop report. Acta Neurochir (Wien). 2017; 159(7):1187–1195.

20. Krieg SM, Shiban E, Buchmann N, et al. Utility of presurgical navigated transcranial magnetic brain stimulation for the resection of tumors in eloquent motor areas. J Neurosurg. 2012; 116(5):994–1001.

21. Retif P, Djibo Sidikou A, Mathis C, et al. Evaluation of the ability of the Brainlab Elements Cranial Distortion Correction algorithm to correct clinically relevant MRI distortions for cranial SRT. Strahlenther Onkol. 2022; 198(10):907–918.

22. Calvo-Ortega JF, Mateos J, Alberich Á, et al. Evaluation of a novel software application for magnetic resonance distortion correction in cranial stereotactic radiosurgery. Med Dosim. 2019; 44(2):136–143.

23. Negwer C, Hiepe P, Meyer B, Krieg SM. Elastic Fusion Enables Fusion of Intraoperative Magnetic Resonance Imaging Data with Preoperative Neuronavigation Data. World Neurosurg. 2020; 142:e223–e228.

24. Gerhardt J, Sollmann N, Hiepe P, et al. Retrospective distortion correction of diffusion tensor imaging data by semi-elastic image fusion - Evaluation by means of anatomical landmarks. Clin Neurol Neurosurg. 2019; 183:105387.

25. Frey D, Strack V, Wiener E, Jussen D, Vajkoczy P, Picht T. A new approach for corticospinal tract reconstruction based on navigated transcranial stimulation and standardized fractional anisotropy values. Neuroimage. 2012; 62(3):1600–1609.

26. Yeh FC, Panesar S, Fernandes D, et al. Population-averaged atlas of the macroscale human structural connectome and its network topology. Neuroimage. 2018; 178:57–68.

27. Wasserthal J, Neher P, Maier-Hein KH. TractSeg - Fast and accurate white matter tract segmentation. Neuroimage. 2018; 183:239–253.

28. Avants BB, Tustison NJ, Song G, Cook PA, Klein A, Gee JC. A reproducible evaluation of ANTs similarity metric performance in brain image registration. Neuroimage. 2011; 54(3):2033–2044.

29. Henker C, Kriesen T, Glass Ä, Schneider B, Piek J. Volumetric quantification of glioblastoma: experiences with different measurement techniques and impact on survival. J Neurooncol. 2017; 135(2):391–402.

30. Sawaya R, Hammoud M, Schoppa D, et al. Neurosurgical outcomes in a modern series of 400 craniotomies for treatment of parenchymal tumors. Neurosurgery. 1998; 42(5):1044–1055; discussion 1055-1046.

31. Lu J, Zhao Z, Zhang J, et al. Functional maps of direct electrical stimulation-induced speech arrest and anomia: a multicentre retrospective study. Brain. 2021; 144(8):2541–2553.

32. Muir M, Prinsloo S, Traylor JI, et al. Transcranial magnetic stimulation tractography and the facilitation of gross total resection in a patient with a motor eloquent glioblastoma: illustrative case. J Neurosurg Case Lessons. 2022; 3(20).

33. Chang WH, Wei KC, Chen PY, et al. The impact of patient factors and tumor characteristics on language neuroplasticity in left hemispheric diffuse gliomas prior to surgical resection. J Neurooncol. 2023; 163(1):95–104.

34. Chang EF, Raygor KP, Berger MS. Contemporary model of language organization: an overview for neurosurgeons. J Neurosurg. 2015; 122(2):250–261.

35. Forkel SJ, Hagoort P. Redefining language networks: connectivity beyond localised regions. Brain Struct Funct. 2024.

36. Tuncer MS, Salvati LF, Grittner U, et al. Towards a tractography-based risk stratification model for language area associated gliomas. Neuroimage Clin. 2021; 29:102541.

37. Corbetta M, Ramsey L, Callejas A, et al. Common behavioral clusters and subcortical anatomy in stroke. Neuron. 2015; 85(5):927–941.

38. Griffis JC, Nenert R, Allendorfer JB, Szaflarski JP. Damage to white matter bottlenecks contributes to language impairments after left hemispheric stroke. Neuroimage Clin. 2017; 14:552–565.

39. Duffau H. A two-level model of interindividual anatomo-functional variability of the brain and its implications for neurosurgery. Cortex. 2017; 86:303–313.

40. Duffau H. Does post-lesional subcortical plasticity exist in the human brain? Neurosci Res. 2009; 65(2):131–135.

41. Talozzi L, Forkel SJ, Pacella V, et al. Latent disconnectome prediction of long-term cognitive-behavioural symptoms in stroke. Brain. 2023; 146(5):1963–1978.

42. Sarubbo S, De Benedictis A, Merler S, et al. Towards a functional atlas of human white matter. Hum Brain Mapp. 2015; 36(8):3117–3136.

43. Muir M, Prinsloo S, Michener H, et al. TMS Seeded Diffusion Tensor Imaging Tractography Predicts Permanent Neurological Deficits. Cancers (Basel). 2022; 14(2).

44. Muir M, Prinsloo S, Michener H, et al. Transcranial magnetic stimulation (TMS) seeded tractography provides superior prediction of eloquence compared to anatomic seeded tractography. Neurooncol Adv. 2022; 4(1):vdac126.

45. Muir M, Gadot R, Prinsloo S, et al. Comparative study of preoperative functional imaging combined with tractography for prediction of iatrogenic motor deficits. J Neurosurg. 2023; 139(1):65–72.

46. Scholz J, Klein MC, Behrens TE, Johansen-Berg H. Training induces changes in white-matter architecture. Nat Neurosci. 2009; 12(11):1370–1371.

47. Roberts TP, Liu F, Kassner A, Mori S, Guha A. Fiber density index correlates with reduced fractional anisotropy in white matter of patients with glioblastoma. AJNR Am J Neuroradiol. 2005; 26(9):2183–2186.

48. Sternberg EJ, Lipton ML, Burns J. Utility of diffusion tensor imaging in evaluation of the peritumoral region in patients with primary and metastatic brain tumors. AJNR Am J Neuroradiol. 2014; 35(3):439–444.

49. Basser PJ, Mattiello J, LeBihan D. MR diffusion tensor spectroscopy and imaging. Biophys J. 1994; 66(1):259–267.

50. Beaulieu C. The basis of anisotropic water diffusion in the nervous system - a technical review. NMR Biomed. 2002; 15(7-8):435–455.

51. Sarubbo S, Tate M, De Benedictis A, et al. Mapping critical cortical hubs and white matter pathways by direct electrical stimulation: an original functional atlas of the human brain. Neuroimage. 2020; 205:116237.

52. Roberts DW, Hartov A, Kennedy FE, Miga MI, Paulsen KD. Intraoperative brain shift and deformation: a quantitative analysis of cortical displacement in 28 cases. Neurosurgery. 1998; 43(4):749–758; discussion 758-760.

53. Hartkens T, Hill DL, Castellano-Smith AD, et al. Measurement and analysis of brain deformation during neurosurgery. IEEE Trans Med Imaging. 2003; 22(1):82–92.

54. Ille S, Schwendner M, Zhang W, Schroeder A, Meyer B, Krieg SM. Tractography for Subcortical Resection of Gliomas Is Highly Accurate for Motor and Language Function: ioMRI-Based Elastic Fusion Disproves the Severity of Brain Shift. Cancers (Basel). 2021; 13(8).

55. Elias WJ, Fu KM, Frysinger RC. Cortical and subcortical brain shift during stereotactic procedures. J Neurosurg. 2007; 107(5):983–988.

56. Petersen EA, Holl EM, Martinez-Torres I, et al. Minimizing brain shift in stereotactic functional neurosurgery. Neurosurgery. 2010; 67(3 Suppl Operative):ons213-221; discussion ons221.

57. Zhang W, Ille S, Schwendner M, Wiestler B, Meyer B, Krieg SM. Tracking motor and language eloquent white matter pathways with intraoperative fiber tracking versus preoperative tractography adjusted by intraoperative MRI-based elastic fusion. J Neurosurg. 2022; 137(4):1114–1123.

58. Ille S, Schroeder A, Wagner A, et al. Intraoperative MRI-based elastic fusion for anatomically accurate tractography of the corticospinal tract: correlation with intraoperative neuromonitoring and clinical status. Neurosurg Focus. 2021; 50(1):E9.

59. Duffau H, Thiebaut de Schotten M, Mandonnet E. White matter functional connectivity as an additional landmark for dominant temporal lobectomy. J Neurol Neurosurg Psychiatry. 2008; 79(5):492–495.

60. Matarrese MAG, Loppini A, Fabbri L, et al. Spike propagation mapping reveals effective connectivity and predicts surgical outcome in epilepsy. Brain. 2023; 146(9):3898–3912.

61. Tournier JD, Calamante F, Connelly A. MRtrix: diffusion tractography in crossing fiber regions. International journal of imaging systems and technology. 2012; 22(1):53–66.

